# SeuratIntegrate: an R package to facilitate the use of integration methods with Seurat

**DOI:** 10.1101/2024.12.16.628691

**Authors:** Florian Specque, Aurélien Barré, Macha Nikolski, Domitille Chalopin

## Abstract

**Motivation:** Integrating multiple datasets has become an increasingly common task in scRNA-seq analysis. The advent of single-cell atlases adds further complexity to this task, as they often involve combining data with complex, nested batch effects - such as those arising from multiple studies, organs or disease states. Accurate data integration is essential to distinguish cell types with sufficient granularity, thereby reflecting true biological patterns, and to create reliable reference datasets for the community. In this context, the latest version of Seurat (v5) introduced a multi-layered object structure to facilitate the integration of scRNA-seq datasets in a unified manner. However, the panel of available batch-correction methods remains limited to five algorithms within Seurat, restricting users from accessing a broader diversity of available tools, particularly Python-based methods. Furthermore, no existing R tool assists the user in making an informed decision in selecting the most appropriate integration approach.

**Results:** To overcome these challenges, we developed SeuratIntegrate, an open source R package that extends Seurat’s functionality. SeuratIntegrate supports eight integration methods, incorporating both R- and Python-based tools, and enables performance evaluation of integration through several scoring methods. This functionality allows for a more versatile and informed integration process.

**Availability:** SeuratIntegrate is available at https://github.com/cbib/Seurat-Integrate/. The package is released under the MIT License.

## 1. INTRODUCTION

The rapid growth of single-cell datasets in recent years has enabled the possibility to study multiple samples from different biological and technical contexts, all aggregated in large atlases. These atlases compile data from different experiments, reagent batches or sequencing technologies. While these resources are invaluable for advancing single-cell based studies, bringing together such large datasets presents the challenge of overcoming confounding effects that may mask true biological differences and thus potentially impact analyses such as for example differential expression (Argelaguet et al. 2021; Nguyen et al. 2023; Maan et al. 2024).

Seurat v5 (Hao et al. 2024) and Scanpy (Wolf et al. 2018) are the most widely used R-based and Python-based tools, respectively, for single-cell data analysis. Both tools offer comprehensive functionalities, including visualization, clustering, and trajectory analysis. A key feature of both is their support for batch-effect correction and data integration, essential steps for harmonizing datasets from diverse biological and technical contexts.

Batch-effect correction and integration are embedded within these tools, but they differ in the methods they provide. Seurat utilizes an ‘anchor-based’ strategy for integration based on Mutual Nearest Neighbors (MNN) for batch-effect correction. V5 added native support for Harmony and scVI. Scanpy incorporates a variety of methods, including BBKNN, ComBat, Scanorama, Harmony, and several deep-learning algorithms, offering flexibility in addressing batch effects and aligning datasets. Notably, some of these methods, implemented in Seurat and Scanpy, were included in the benchmarking study by Luecken et al. (2020), underscoring their relevance and widespread use.

Luecken et al. (2020) evaluated the performance, usability, and scalability of over 20 integration and batch-effect correction methods. Their findings demonstrated that the effectiveness of these methods depends on data complexity, such as the number of samples, cell type diversity, and technology differences. Importantly, Python-based methods often outperformed R-based methods on complex real-world datasets. These results emphasize the value of combining Python- and R-based approaches to create a flexible and comprehensive toolkit for single-cell data analysis.

Here, we present ‘SeuratIntegrate’, a R package designed as an extension of Seurat allowing a seamless use of integration methods written either in R or in Python, thereby simplifying cross-platform interoperability. To date, SeuratIntegrate proposes 3 R-based methods and 5 Python-based methods as well as additional functions for performance evaluation.

## 2. Package description and implementation

### 2.1 Integration Methods provided by SeuratIntegrate

SeuratIntegrate extends the functionality of Seurat v5 by providing access to additional integration methods not included in the original package, particularly those written in Python. It supports eight complementary methods for batch-effect correction and data integration, as well as ten metrics to evaluate their efficacy (Figure 1a). These metrics include measures of batch-effect removal and biological signal preservation, allowing for a comprehensive assessment of each method (see section 2.2 for details). This makes it possible to perform robust comparisons across a diverse range of integration techniques.

**Figure 1.**
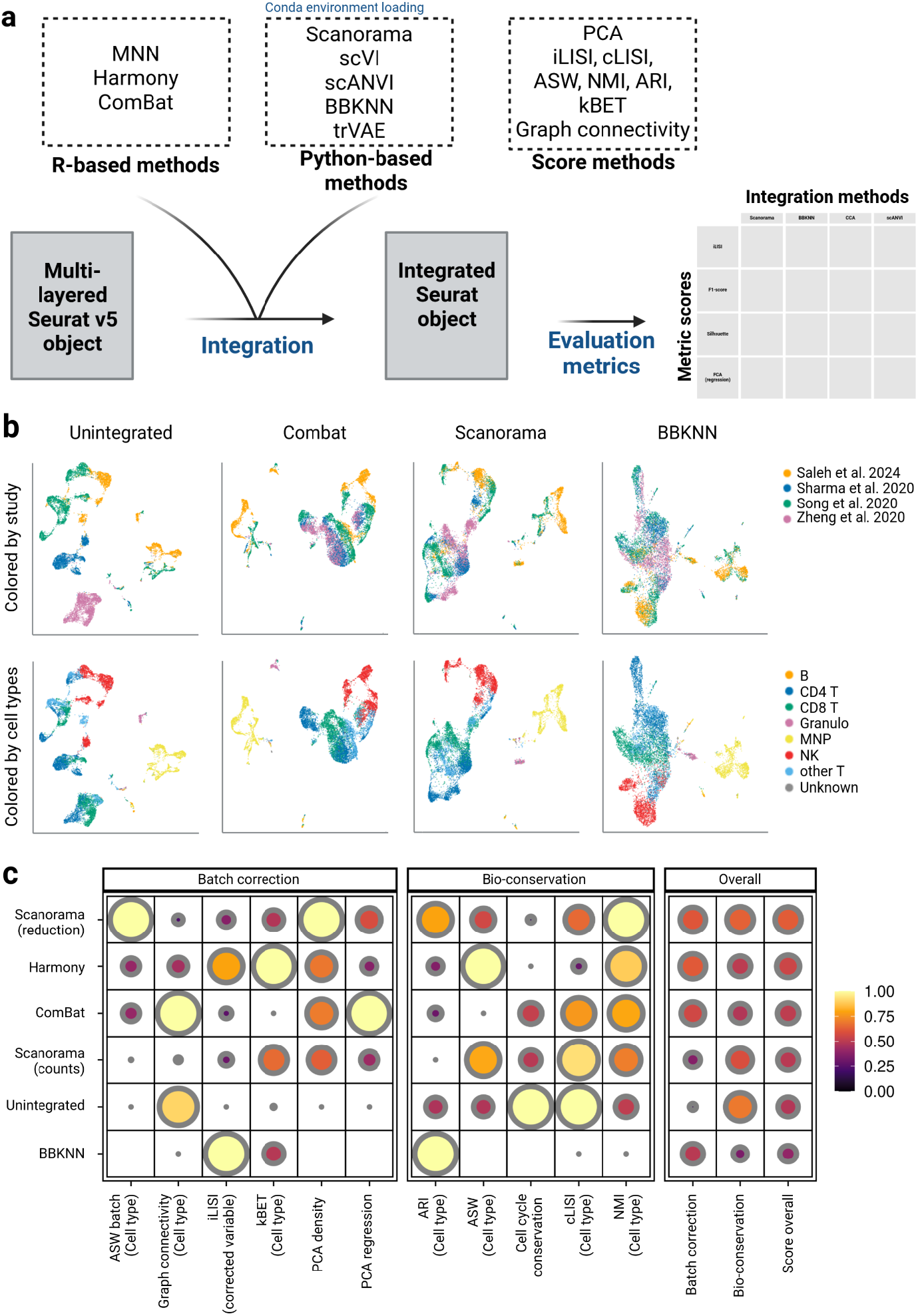
Overview of SeuratIntegrate workflow.

#### Supported Integration Methods

SeuratIntegrate offers both R- and Python-based methods, enabling users to harness a wide range of computational approaches for dataset harmonization (see Table 1).

**Table 1.**
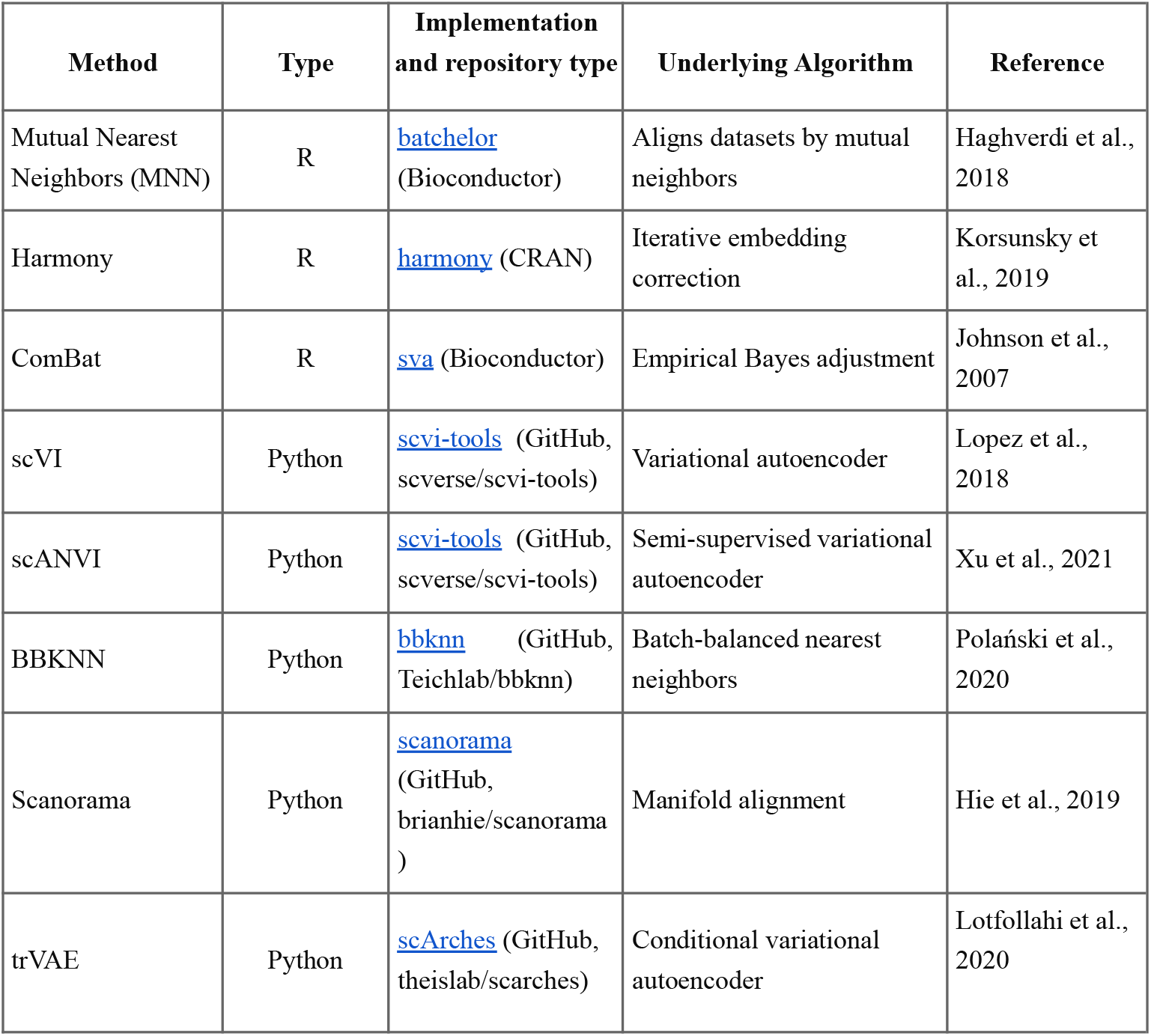
Comprehensive overview of the integration methods provided bySeuratIntegrate. For each method, the following details are provided: original implementation language (R-or Python-based), link to the GitHub repository of this initial implementation, a brief description of the basic algorithm and reference to the original paper where the method was introduced.

#### Key Functionality: *DoIntegrate*

Building on Seurat’s `IntegrateLayers` function introduced in version 5, SeuratIntegrate implements a new function called `DoIntegrate`. It provides users with:

- The ability to launch multiple integrations in a single command,
- Customizable parameters for each integration method,
- Flexibility in choosing the input matrix type and the features to include.

As highlighted by Luecken et al. (2022), the choice of input matrix can have a significant impact on integration performance. By allowing users to select the most appropriate matrix (raw, normalized or scaled) and features, `DoIntegrate`ensures flexibility in the use of integration workflows. Additionally, the function is compatible with other Seurat-supported integration methods, such as CCA, RPCA, and FastMNN (available in SeuratWrappers), enabling seamless interoperability across different integration approaches.

#### Reimplementation of Harmony and scVI

Although Harmony and scVI are already supported by Seurat, they have been re-implemented in SeuratIntegrate to provide users with enhanced flexibility:

- Harmony: Implemented via the `RunHarmon`function, adhering to the original developers’ guidelines. A seed is set by default to ensure reproducible results.
- scVI: Allows users to fine-tune specific parameters, such as the number of layers in the neural network and their size, providing greater control over the integration process.

##### Python integration via reticulate

SeuratIntegrate leverages the `reticulate` package (Ushey et al. 2024) to enable Python-based integration methods. To streamline this process, the package includes tools to create and manage Conda environments for each Python-based method. However, since R sessions can only load one Python environment at a time, SeuratIntegrate overcomes this limitation by using the `Future` package (Bengtsson 2020) to launch background R sessions for each environment. This functionality is seamlessly integrated into the `DoIntegrate` function.

For more details on using these features and additional functionalities, such as connectivity computation from cell-to-cell distances, please refer to the GitHub vignettes.

#### 2.2 Evaluation metrics Metrics

SeuratIntegrate provides a suite of evaluation metrics to compare integration results, many of which have been previously described in Luecken et al. (2022) and Tran et al. (2020). These metrics include Local Inverse Simpson’s Index (LISI) comprising iLISI and cLISI, kBET, PCA-based scores, Average Silhouette Width (ASW), graph connectivity, Normalised Mutual information (NMI), and Adjusted Rand Index (ARI) (Figure 1a). Below, we group these metrics based on whether they require known cell type labels or can be used in a label-free manner:

Label-based Metrics:

- cLISI: Measures the preservation of cell type purity.
- NMI and ARI: Focus on classification accuracy.
- Silhouette (ASW): Evaluates within- and between-cluster similarity and can be calculated with or without labels (batch-based).
- Graph Connectivity: Estimates how well cells with the same identity label share neighborhoods in the k-nearest neighbor (KNN) graph.

Label-free Metrics:

- iLISI and kBET: Measure batch mixing using graph-based methods. SeuratIntegrate allows users to adjust the parameter k for these metrics.
- PCA-related Scores:
- RPCA: Quantifies batch removal via principal component regression.
- DPCA: Uses Gaussian kernel density overlap as a proxy for batch removal.
- Cell cycle conservation: Evaluates cell-cycle conservation before and after batch correction.

#### User-Friendly Features

To streamline evaluation, SeuratIntegrate allows users to save multiple scores for different integration methods directly within the Seurat object as a two-dimensional array using functions starting with `AddScore`. All scores can be scaled between 0 and 1 for easier interpretation. Additionally, a batch correction score and a bio-conservation score can be computed as a mean of the corresponding metrics. To ensure a more balanced contribution of each score, min-max rescaling can be performed beforehand. Finally, an overall score, calculated as the weighted sum of batch correction and bio-conservation scores for each method, is provided to guide users in selecting the best-performing integration.

#### Visualization Options

SeuratIntegrate includes three plotting options — dotplot, lollipop, and radar plots — within a unique function `PlotScores` to facilitate visual comparisons of integration performance. These plots are built using ggplot2, ensuring compatibility with popular visualization workflows.

a. Summary of methods available in SeuratIntegrate either for integration or evaluation of the integration.
b. Running example of the use of SeuratIntegrate with eight samples downloaded from four different studies, containing immune cells from liver tumor microenvironment. UMAPs from unintegrated, as well as ComBat, Scanorama (reduction method) and BBKNN integrated data are colored by the origin of the study (upper) or by manual cell type annotation (lower). UMAP from Scanorama (counts-based method) and Harmony are not shown.
c. Dotplot displaying evaluation scores for various integration methods. Absence of a dot indicates metrics that were not evaluated due to the characteristics of the integration method. For example, PCA scores are not computed for BBKNN because it returns a graph instead of numerical components. Label-based metrics were calculated using manually curated annotation. Scores are organized in two main categories: batch correction and biological conservation. In addition to individual metrics, overall scores for batch correction, biological conservation, and total performance are also included.

## 3. USAGE EXAMPLE

### Dataset Preparation

To demonstrate the utility of SeuratIntegrate, we analyzed immune cells from the hepatocellular carcinoma microenvironment. This dataset comprises eight samples from four publicly available studies: Sharma et al. (2020) [GSE156337], Song et al. (2020) [CRA002308], Zheng et al. (2020) [CRA001276], and Giraud et al. (2024) [GSE245909]. All samples were generated using 10x Genomics technology and sequenced on an Illumina platform. After downloading the raw data, we used CellRanger v7.1.0 to generate raw gene count matrices, followed by cell quality filtering, doublet and non-immune cell removal, and merging into a multi-layered Seurat object (∼40,000 cells). For integration, we randomly selected 10,000 cells to speed up the analysis. Scripts to generate the example object as well as the multi-layered object are available on a Zenodo deposit (accession 10.5281/zenodo.14288361).

### Integration

The unintegrated data revealed a clear bias in cell distribution based on the study of origin, as visualized in UMAP plots (Figure 1b ‘Unintegrated’ UMAP). For instance, CD4 T, CD8 T, and other T cells were segregated by study rather than biological cell type, highlighting the impact of batch effects and justifying the need for integration. Indeed, in a correct integration, we would expect cells to intermingle at the study level while remaining distinct by cell type.

Using SeuratIntegrate, we applied five integration methods — ComBat, Scanorama (based on counts or reduction), Harmony, and BBKNN — to address this issue (Figure 1b,c). After integration, UMAP plots showed improved mixing of cells from different studies while preserving cell type separation, demonstrating the effectiveness of these methods in reducing batch effects and aligning datasets.

### Evaluation

To compare integration methods, we used the evaluation metrics implemented in SeuratIntegrate, including batch correction scores and biological knowledge conservation scores. Biological knowledge was assessed at three annotation levels: Azimuth liver level 1, Azimuth liver level 2 (Hao et al. 2021), and a manual annotation containing eight cell types (Figure 1c - only shows label scores from the manual annotation). Scores were saved within the Seurat object using `AddScore` functions, scaled between 0 and 1, normalised using min-max rescaling prior to computing overall scores and visualized using dotplot.

### Findings

The dotplot revealed distinct performance patterns across the different methods (Figure 1c):

- Batch Correction: Harmony outperformed all methods with a score of 0.616. Unsurprisingly, the unintegrated data exhibits the worst score in this regard (0.162). In between, in descending order of performance, we find Scanorama (reduction method), ComBat, BBKNN and finally Scanorama (counts method) with a scaled score of 0.607, 0.56, 0.496 and 0.36, respectively.
- Biological Conservation: The unintegrated data reaches the highest score (0.69), followed by ComBat and Scanorama (counts method) (0.64 each) then Scanorama (reduction method) and BBKNN (0.62 each) and Harmony (0.6)

Several of our observations are in line with the conclusions of Luecken et al. (2022) regarding the human immune dataset. Scanorama (reduction method) presents a higher overall score compared to other methods, followed by Harmony. Scores computed on Scanorama’s outputs are usually higher for the integrated reduction than for the corrected counts.

However, some unexpected results can be noted in the overall ranking. First, the unintegrated data reaches the best biological conservation score, which can be explained by the highest cell cycle conservation and cLISI scores. Indeed, cell cycle conservation is scaled by comparing the cell cycle phase of cells to the unintegrated data. Hence, the unintegrated data always has the maximum score of 1. Furthermore, cLISI only considers the cell type diversity within each cell’s neighborhood, regardless of the batch of origin. In this respect, cells with the same annotation are not clustered together (Figure 1b, lower left). Second, BBKNN is the poorest-performing method in our analysis, which can be attributed to its very low scores for two out of three bio-conservation metrics tested (namely, cLISI and NMI). Indeed, computing min-max normalisation before aggregating overall scores, can amplify score differences, particularly when comparing a small number of methods. Hence, an integration method that performs poorly on two scores is likely to rank among the lowest overall. As a rule of thumb, min-max rescaling is well-suited for comparing a large number of integrations. However, its reliability diminishes as the number of integrations decreases, warranting careful manual evaluation in such cases.

## 4. CONCLUSION

The proposed R package named SeuratIntegrate facilitates the use of integration methods for single-cell transcriptomic data allowing easy interoperability between R and Python. We illustrated how SeuratIntegrate facilitates comprehensive integration and evaluation workflows by applying it to a dataset composed by immune cells from the hepatocellular carcinoma microenvironment. The observed variability in performance underscored the importance of dataset-specific evaluations, as demonstrated by our results and previous studies (e.g., Luecken et al. 2022).

The source code, installation process and vignettes demonstrating usage are freely available on GitHub: https://github.com/cbib/Seurat-Integrate.

